# Increased remodeling and impaired adaption to endurance exercise in desminopathy

**DOI:** 10.1101/2021.10.03.462939

**Authors:** Agata A. Mossakowski, Henning T. Langer, Alec Bizieff, Alec M. Avey, Hermann Zbinden-Foncea, Marc Hellerstein, Keith Baar

**Affiliations:** Neurobiology, Physiology and Behavior, University of California, Davis, CA; Department of Physiology and Membrane Biology, University of California, Davis, CA; Charité – Universitätsmedizin Berlin, corporate member of Freie Universität Berlin, Humboldt-Universität zu Berlin, and Berlin Institute of Health; Department of Nutritional Sciences and Toxicology, University of California, Berkeley, USA; Exercise Physiology and Metabolism Laboratory, School of Kinesiology, Faculty of Medicine, Universidad Finis Terrae, Santiago. Chile

**Keywords:** Muscular Dystrophy, Chronic Injury, Desminopathy, Intermediate filament, Mechanical vulnerability, Protein aggregation myopathy

## Abstract

Desminopathy the most common intermediate filament disease in humans. Desmin is an essential part of the filamentous network that aligns myofibrils, anchors nuclei and mitochondria, and connects the z-discs and the sarcolemma. We created a rat model with a mutation in R349P *DES*, analog to the most frequent R350P *DES* missense mutation in humans. To examine the effects of a chronic, physiological exercise stimulus on desminopathic muscle, we subjected R349P DES rats and their wildtype (WT) and heterozygous littermates to a treadmill running regime. We saw significantly lower running capacity in DES rats that worsened over the course of the study. We found indicators of increased autophagic and proteasome activity with running in DES compared to WT. Stable isotope labeling and LC-MS analysis displayed distinct adaptations of the proteomes of WT and DES animals at baseline as well as with exercise: While key proteins of glycolysis, mitochondria and thick filaments increased their synthetic activity with running in WT, these proteins were higher at baseline in DES and did not change with running. The results suggest an impairment in adaption to chronic exercise in DES muscle and a subsequent exacerbation in the functional and histopathological phenotype.

## INTRODUCTION

Desmin is the most abundant intermediate filament in mature skeletal, cardiac, and smooth muscle. The 53-kDa protein plays an integral role in the structure and alignment of myofibrils and in the resistance of muscle cells to mechanical stress by forming a scaffold that links the Z-disc to the subsarcolemmal cytoskeleton and connects the contractile apparatus with mitochondria, the nucleus, the sarcolemma, the T-tubules and the post-synaptic areas of motor endplates^1^. Over 60 disease-causing mutations that spread over the entire desmin gene on chromosome 2q35 have been described^2^. The resulting desminopathies are a clinically heterogenous group of myofibrillar muscular dystrophies that are commonly associated with skeletal and cardiomyopathy, but can vary widely in their clinical onset, severity, and progression^3–9^.

The exact molecular pathogenesis of desminopathies remains to be fully understood. Most desmin mutations generate a desmin protein incapable of forming the normal filamentous network. Mutant desmin can escape proteolytic degradation and form the insoluble intracellular aggregates typical of all myofibrillar myopathies, which disrupt preexisting filamentous networks and have therefore been described as “toxic”^10,11^. While desmin is expressed very early in the development and differentiation of skeletal and cardiac muscle^12–14^, first symptoms of desminopathy occur mainly between the second and fifth decade of life^6,15^. It is thought that this seeming discrepancy reflects the eventual failure of the protein quality control (PQC) system^16,17^. Once the PQC gets overwhelmed by misfolded proteins and sarcoplasmic aggregates, regenerative mechanisms cannot keep up with the ensuing myofibrillar death, and the loss of functional muscle fibers leads to symptoms of myopathy like progressive muscular weakness and wasting, cardiomyopathy with conduction anomalies and shortness of breath.

Consequently, the main histopathological hallmarks of desminopathy are sarcoplasmic and subsarcolemmal desmin-positive protein aggregates^18–21^. Other myopathic features include increased fiber size variability, central nuclei, atrophic or necrotic fibers, fiber splitting, rimmed vacuoles, inflammation and increased intramuscular connective and fat tissue^18,22,23^.

The most prevalent mutation in humans is caused by the exchange of arginine to proline at position 350 of the desmin gene (R350P)^8,24,25^. Clemen et al. created R349P knock-in mice, ortholog to the human R350P missense mutation, that modeled the human disease both histopathologically and functionally^10^. More recently, we engineered a CRISPR-Cas9 knock-in rat with the same mutation to create a model organism that is physiologically closer to humans. While we saw a clear histopathological phenotype with sarcoplasmic aggregates and signs of regeneration in the muscle of 6-month-old mutants, muscle mass and force did not differ significantly from wildtype littermates even after synergist ablation^26^. The observed histopathological and molecular indicators of injury and regeneration were exacerbated by an acute eccentric contraction challenge in older animals^27^. However, in general we found our model to result in a relatively mild skeletal muscle phenotype and hypothesized that chronic loading was needed to trigger the pathophysiology seen in the human condition. To provide an adequate stimulus to trigger the pathophysiology, we subjected older rats to a 6-week long treadmill running regime that included 4 weeks of downhill running and then studied the running performance, histology, biochemistry, and protein turnover in the wild type, heterozygous, and homozygous mutant rats.

## RESULTS

### Body composition and muscle weights

Rats were subjected to 6 weeks of running (Fig1A). DES rats weighed less than their WT and HET littermates at an average of 564±63 g (DES) compared to 637±32 g (WT) and 657±71 g (HET) (Fig1B) (p=0,02 for genotype, p=0.31 for running and p=0.92 for interaction).

**Figure 1 -.**
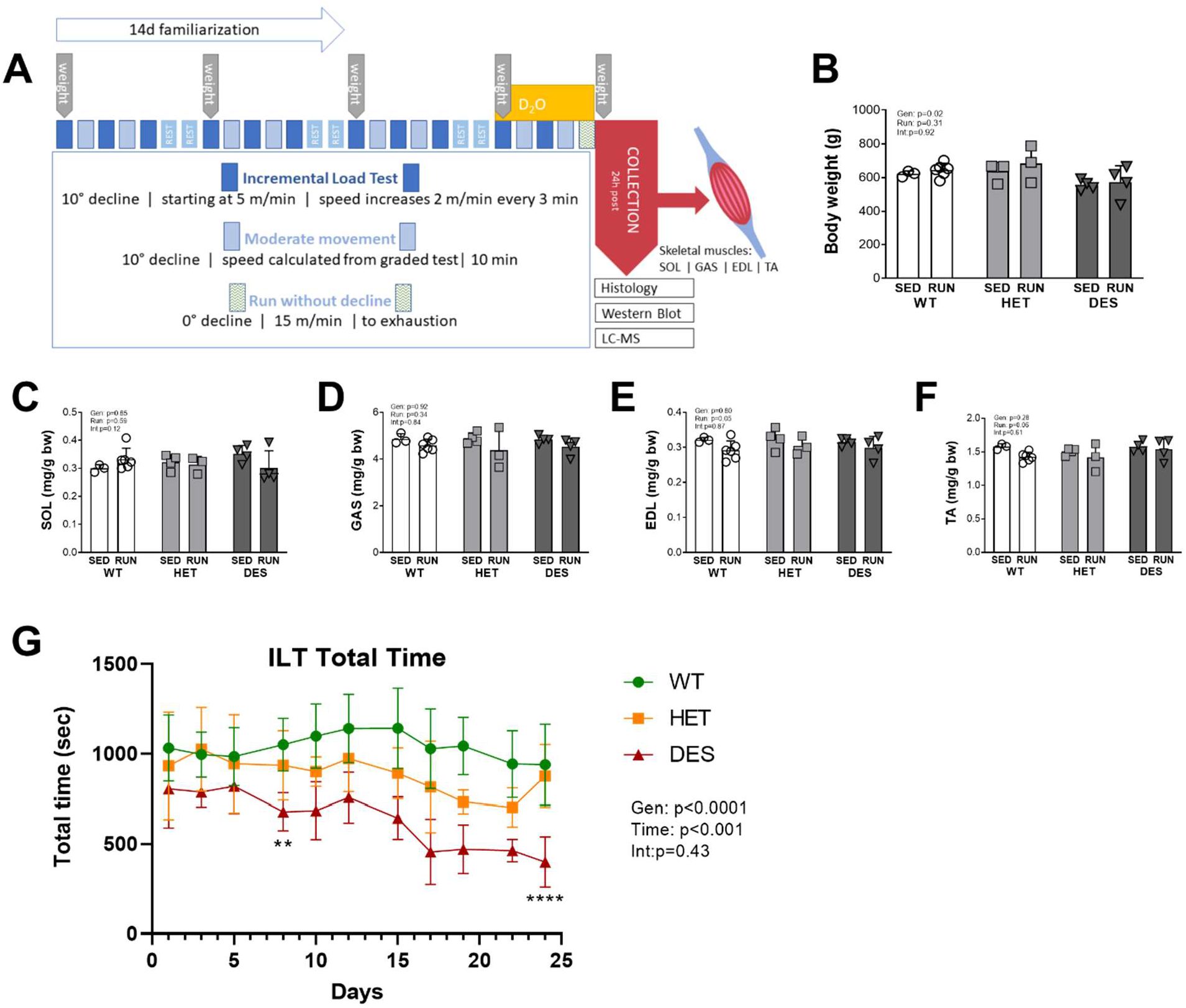
Running protocol and muscle collections. Rats in the exercise group were subjected to downhill running at a decline of −10° for four weeks after two weeks of treadmill familiarization. Rats performed an incremental load test (ILT) on day 1, 3 and 5 of each week. Treadmill speed was started of 5 m/min and increased by 2 m/min every 3 min until exhaustion. Endogenous deuterium levels were raised to about 4 % by a priming intraperitoneal injection of 0.014 mL *g−1 bodyweight deuterium oxide in 0.9 % NaCl five days before collection. Deuterium (D2O) levels were maintained via drinking water. (A). DES rats (n=8) had lower body weight than WT (n=9) and HETs (n=7) (B). Muscles were collected 24h after the last run. Muscle weights are presented relative to body weight (C-F). SOL = soleus, EDL = extensor digitorum longus, GAS = gastrocnemius, TA = tibialis anterior muscle (C-F). Results for mean total running time at each ITL are presented in (G). Gen=Genotype, Run=Running, Int=Interaction, Time=effect of days of training; AU=arbitrary units; Sed=sedentary controls. Group differences were assessed via a two-way ANOVA, error bars indicate ± SD. ** indicates p < 0.01 and **** indicates p < 0.0001

Absolute muscle weights were also lower in DES rats in all collected muscles, but not when calculated relative to body weight (Fig1C-F). Contrary to our expectation, muscle weights relative to body weight were decreased in the running group compared to sedentary controls in gastrocnemius (GAS) (Fig1D, p=0.092 for genotype, p=0.03 for running and p=0.84 for interaction) and extensor digitorum longus (EDL) muscle (Fig1E, p=0.8 for genotype, p=0.05 for running and p=0.87 for interaction), and the same trend was found in tibialis anterior (TA) muscle (Fig1F, p=0.29 for genotype, p=0.06 for running and p=0.61 for interaction) but here it did not reach statistical significance. In soleus (SOL), muscle weight tended to go up in the running group in WT and down in DES, but again without reaching statistical significance (Fig1C, p=0.85 for genotype, p=0.59 for running and p=0.12 for interaction).

One animal (DES) had to be terminated due to weight loss that met humane endpoint criteria in week 2. Post-mortem examination showed a severe bowel obstruction.

### Chronic exercise exacerbates a functional phenotype

The running protocol was designed to induce moderate chronic injury of the loaded muscles. In the first incremental load test (ILT), WT rats ran for a mean of 1032±168 s, HET rats for 933±244 s and DES rats for 804±188 s. The WT and HET animals maintained their maximal running time, whereas time to exhaustion decreased continuously in DES rats. In the last ILT, WT rats ran for a mean of 940±206 s, HET rats for 876±143 s and DES rats for 399±122 s (Fig1G). Overall, performance in all genotypes decreased during the course of the experiment – WT by 9% (most of this decrease stemming from one animal that dropped disproportionally in performance over the last two ILT), HET by 6% and DES by 50% (p<0.0001 for genotype, p<0.001 for time and p=0.43 for interaction). Running time declined markedly earlier in DES (first decline of >5% on ILT4, p<0.01 with Tukey’s multiple comparisons test) than in HET (first decline of >5% on ILT8, p=0.22 with Tukey’s multiple comparisons test) and WT (first decline of >5% on ILT10, p=0.74 with Tukey’s multiple comparisons test) (Fig1G).

### Histopathological changes show signs of ongoing myopathy in DES animals

Histopathological changes typical of myopathies, such as central nuclei, atrophic and necrotic fibers, and fiber size variability were prominent in H&E-stained cross-sections of sedentary DES SOL and were moderately aggravated in running DES SOL (Fig2A). HET SOL appeared predominantly normal in H&E, but had a slightly increased number of myofibers with central nuclei compared to WT. While sedentary WT showed an average of 0.8±0.06% of fibers (15±1 absolute count) with central nuclei, 2±0.7% of HET myofibers (33±16 absolute count) had central nuclei, which was noticeable in a qualitative analysis, but not statistically significant (p=1 with Tukey’s multiple comparisons test). The percentage of fibers with central nuclei was significantly higher in DES 18±5% (475±73 absolute count) (p<0.0001 for genotype, p=0.21 for running and p=0.05 for interaction). While the ratio of fibers with central nuclei was similar in the sedentary and running group of WT and HET (both p=1 with Tukey’s multiple comparisons test), the DES running group trended to have a lower percentage of fibers with central nuclei at 15±6% (315±132 absolute count, p=0.07 with Tukey’s multiple comparisons test) compared to the sedentary DES rats (Fig2B). Whole muscle cross sectional analysis showed significantly smaller fibers in sedentary DES rats at a mean minimal Feret’s diameter of 46±1 µm compared to sedentary WT (63±1 µm) and HET (53±5 µm). Fiber size was not significantly impacted by running for any of the genotypes (p<0.0001 for genotype, p=0.23 for running and p=0.11 for interaction) (Fig2C). Frequency distribution analysis of Feret’s diameter in whole muscle sections reveals a shift towards smaller fibers and a broader curve, representative of fiber size variability in DES (Fig2G). Changes in fiber size did not correlate with the change in performance between the first and last ILT (R^2^=0.22, data not shown). Fiber number tended to increase with running in WT (1875±298 in sedentary vs 2520±339 in running) while it tended to decrease with running in DES (2582±243 in sedentary vs 2345±416 in running) (p=0.27 for genotype, p=0.3 for running and p=0.06 for interaction) (Fig2E).

**Figure 2 -.**
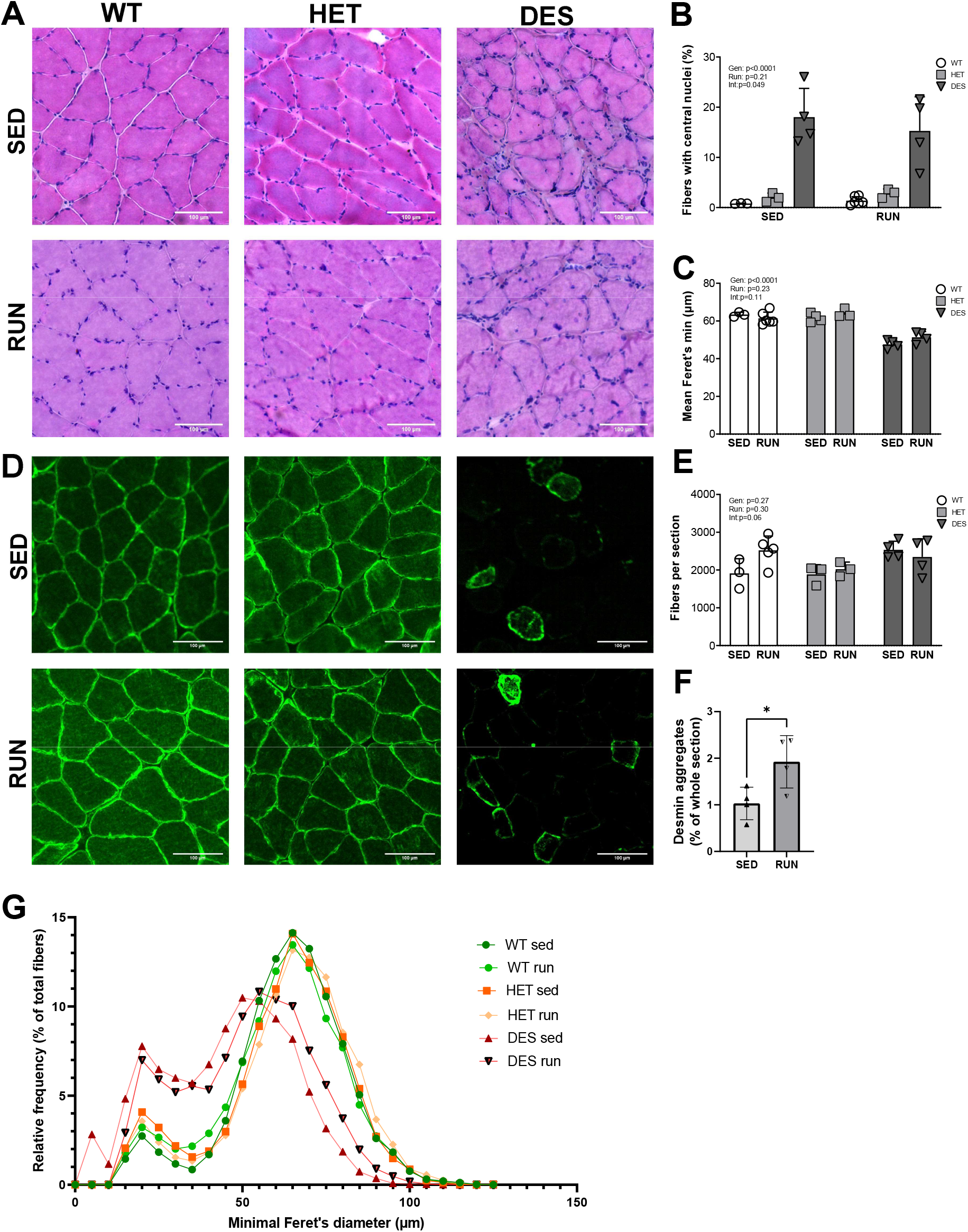
Histological changes in DES mutated muscle. H&E staining of soleus muscle (SOL) sections showed an increased number of necrotic and regenerating fibers (A) and an increased number of fibers with central nuclei (A&B) in desmin mutants. Immunohistochemical staining for desmin (green) showed distribution around the sarcolemma of WT and HET fibers, but only aggregates in DES animals (C) Desmin aggregates were quantified in relation to whole muscle section (F). The mean minimal Feret’s diameter calculated from all fibers of SOL cross sections was lower in DES compared to WT (C), and relative frequency analysis of all fibers showed a shift to a higher number of smaller fibers in DES (G). Fiber number tended to increase with running in WT, while it tended to decrease with running in DES (E). Gen=Genotype, Run=Running, Int=Interaction; AU=arbitrary units; Sed=sedentary controls. (*n* for WT sed=3, WT run=6, HET sed=4, HET run=3, DES sed=4, DES run=4). Group differences were assessed via a two-way ANOVA, error bars indicate ± SD. * indicates p ≤ 0.05

Immunohistochemical analysis of desmin showed subsarcolemmal and cytosolic desmin-positive aggregates in DES animals, a classic hallmark of desminopathy (Fig2D). Desmin aggregates relative to whole section area were higher in the running group (1.92±0.49 %) compared to the sedentary group (1.03±0.3 %, p=0.04) (unpaired t-test, Fig2F).

The integrity of the sarcolemma was assessed by quantifying the number of fibers that stained positive for mouse-IgG. No myofibers of the sedentary animals showed signs of membrane damage, and only one DES animal had 0.13% IgG-positive fibers in the running cohort, while all other sections were negative for IgG (data not shown).

### Cytoskeleton, membrane repair and metabolism

As expected, biochemical analysis of desmin protein levels demonstrated lower desmin in sedentary HETs (46%) and DES mutant animals (69%) compared to sedentary WT (Fig3A). Running decreased desmin protein levels by 43% compared to the sedentary condition in WT rats. However, desmin protein levels increased by 51% in HETs and by 68% in DES compared to the sedentary animals (p for genotype <0.01, p for running = 0.83, p for interaction <0.01, Fig3A). Dysferlin, a protein mainly associated with membrane repair and fusion of repair vesicles to the plasma membrane, tended to be higher in DES rats. There was no effect of running or an interaction between running and the genotype (p for genotype =0.06, p for running = 0.33, p for interaction = 0.52, Fig3B). Annexin A2, which interacts with dysferlin during membrane repair, was not different between genotypes, but increased with running (p for genotype =0.96, p for running < 0.01, p for interaction = 0.66, Fig3C). Muscle LIM (mLIM), a protein prominent in the establishment and maintenance of the myocyte cytoskeleton, was on average 1.4-fold higher in the HET and 2.2-fold higher in the DES compared to the WT rats in sedentary animals. Running increased mLIM levels in all genotypes, but while mLIM increased 2.1-fold in WT and 1.7-fold in HET, DES only increased marginally by 1.03-fold (p for genotype < 0.001, p for running < 0.0001, p for interaction < 0.01, Fig3D). Septin 1 levels were elevated in sedentary DES compared to WT and went down with running. Septin 1 was unchanged in running WT and HET animals (p for genotype = 0.02, p for running = 0.02, p for interaction < 0.01, Fig3E). Dystrophin levels did not differ significantly (p for genotype = 0.25, p for running = 0.85 and p for interaction = 0.63, Fig3F).

**Figure 3 -.**
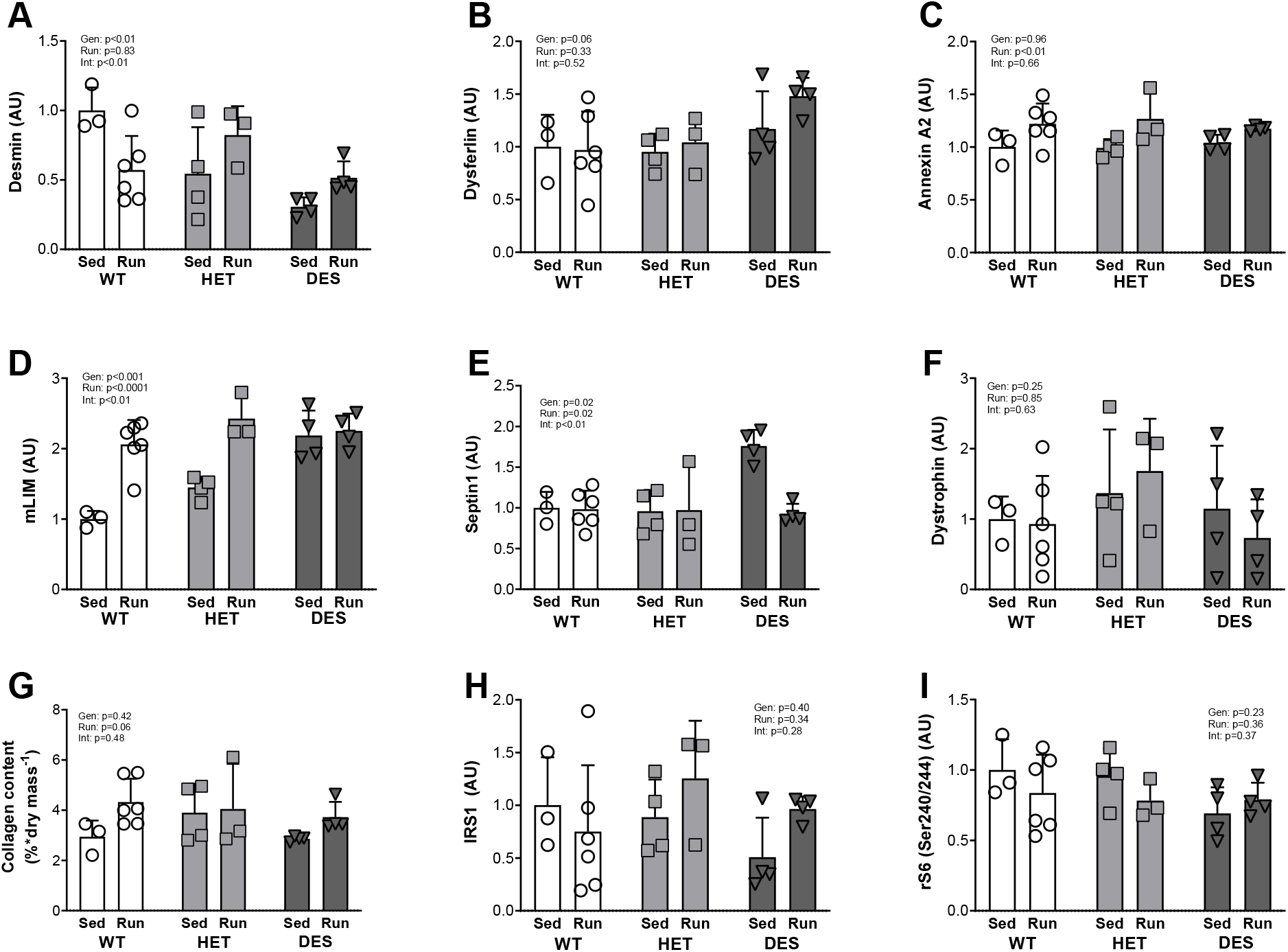
Changes to proteins involved in structural integrity, membrane repair and metabolism. Western Blots demonstrated lower desmin protein levels in sedentary HET (46%) and DES mutant animals (69%) compared to sedentary WT. Desmin levels were decreased with running by 43% compared to sedentary WT rats, but increased by 51% in HET and by 68% in DES compared to the respective sedentary group (A). Dysferlin tended to be higher in DES rats (B). Annexin A2 did not differ between genotypes, but increased with running (C). Muscle LIM (mLIM) in sedentary animals was 1.4-fold higher in the HET and 2.2-fold higher in the DES compared to the WT rats. Running increased mLIM levels in all genotypes (D). Septin 1 was elevated in sedentary DES compared to WT and went down with running. Septin 1 was unchanged in running WT and HET animals (E). Dystrophin levels did not differ significantly (F). Collagen content of whole muscle lysates as assessed by hydroxyproline assay tended to increase after running in all genotypes (G). Insulin receptor substrate 1 (IRS1) (H) and ribosomal protein S6 (rS6 Ser240/244) (I) levels did not differ between groups. Gen=Genotype, Run=Running, Int=Interaction; AU=arbitrary units; Sed=sedentary controls. (n for WT sed=3, WT run=6, HET sed=4, HET run=3, DES sed=4, DES run=4). Group differences were assessed via a two-way ANOVA, error bars indicate ± SD.

Collagen content of whole muscle lysates also did not differ significantly between the groups, although we saw a tendency of collagen levels to increase after running in all genotypes (p for genotype = 0.42, p for running = 0.06, p for interaction = 0.48, Fig3G). Insulin receptor substrate 1 (IRS1) did not differ between groups (p for genotype = 0.40, p for running = 0.34, p for interaction = 0.28, Fig3H). Neither genotype nor running had a significant effect on phosphorylated ribosomal protein S6 (rS6 Ser240/244) levels (p for genotype = 0.23, p for running = 0.36, p for interaction = 0.37, Fig3I).

### Cell damage, degradation, and apoptosis

Next, we investigated proteins involved in the PQC. Upstream autophagy signaling through phosphorylated ULK1 (Ser757) was not affected by genotype but went down with running (p for genotype =0.10, p for running = 0.05, p for interaction = 0.37, Fig4A). Phosphorylated ULK1 (Ser555) tended to be lower in DES and went down with running in all genotypes (p for genotype = 0.15, p for running = 0.04, p for interaction = 0.53, Fig4B). Autophagy related 7 (Atg7), a protein associated with the ubiquitin-proteasome system, decreased slightly in the WT group with running, while its levels increased 2.2-fold in DES animals (p for genotype = 0.01, p for running <0.01 and p for interaction <0.0001, Fig4C). The ratio of LC3 II to LC3 I protein levels was not statistically different between the genotypes and did not change with running (p for genotype = 0.51, p for running = 0.87, p for interaction = 0.68, Fig4D). Heat shock protein 90 (HSP90) differed between the genotypes, whereas running had no effect and there was no interaction between genotype and running (p for genotype = 0.04, p for running = 0.56, p for interaction = 0.41, Fig4E). Heat shock protein 27 (HSP27) levels were the same for all genotypes irrespective of running (p for genotype = 0.26, p for running = 0.58 and p for interaction = 0.92, Fig4F). Free ubiquitin levels went up with running in all groups and tended to increase more in DES than in WT (p for genotype = 0.08, p for running < 0.001, p for interaction = 0.14, Fig4G). The ubiquitination of total proteins, however, went down with running in WT by 23%, stayed the same in HETs and increased by 26% in DES compared to the respective sedentary groups (p for genotype = 0.86, p for running = 0.53, p for interaction <0.01, Fig4H). PTEN-induced kinase 1 (PINK1), a protein involved in autophagy of damaged mitochondria, did not differ significantly between the groups (p for genotype = 0.49, p for running = 0.92 and p for interaction = 0.41, Fig4I). Levels of the autophagy substrate p62 were similar for all genotypes in the sedentary group. Running increased p62 levels in all groups (p for genotype = 0.25, p for running < 0.01, p for interaction = 0.12, Fig4J). Levels of Caspase 3, a cysteine protease that is associated with apoptosis, were significantly higher in the running groups, and elevated by 1.6-fold in WT, 1.7-fold in HET and 2-fold in DES running animals compared to sedentary controls (p for genotype = 0.26, p for running < 0.0001 and p for interaction = 0.42, Fig4K). NOS (pan) levels were reduced 1.6-fold in WT and 1.4-fold in HETs after running, while NOS levels in DES stayed the same after running (p for genotype = 0.09, p for running < 0.001 and p for interaction = 0.12, Fig4L). We observed a similar effect in total OxPhos-levels (p for genotype = 0.91, p for running = 0.04 and p for interaction = 0.44, Fig4M).

**Figure 4 –.**
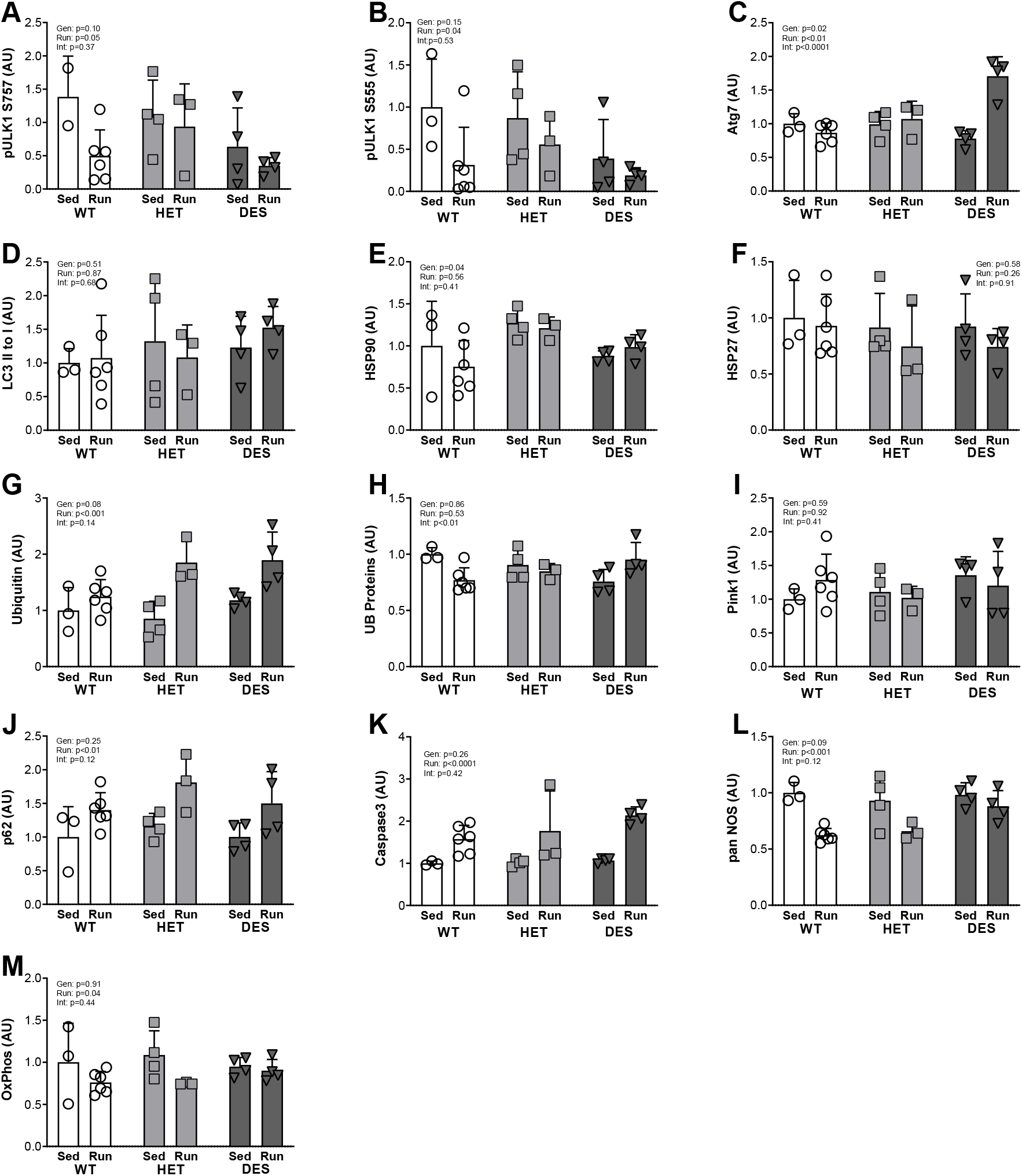
Changes in the Protein Quality Control system. Western Blots of phosphorylated ULK1 (Ser757) showed decreased protein levels in the running group (A). Phosphorylated ULK1 (Ser555) tended to be lower in DES and went down with running in all genotypes (B) Autophagy related 7 (Atg7) decreased in the WT group with running, but increased 2.2-fold in DES animals (C). The ratio of LC3 II to LC3 I protein levels was not statistically different between the genotypes and did not change with running (D). Running had no effect on heat shock protein 90 (HSP90) (E). Heat shock protein 27 (HSP27) levels were the same for all genotypes irrespective of running (F). Free ubiquitin (G), p62 (J) and caspase 3 (K) levels increased with running in all groups. The ubiquitination of total proteins went down with running in WT by 23%, stayed the same in HETs and increased by 26% in DES compared to the respective sedentary groups (H). PTEN-induced kinase 1 (PINK1) did not differ significantly between the groups (I). NOS (pan) levels were reduced 1.6-fold in WT and 1.4-fold in HETs after running, while NOS levels in DES stayed the same after running (L). We observed a similar effect in total OxPhos-levels (M). Gen=Genotype, Run=Running, Int=Interaction; AU=arbitrary units; Sed=sedentary controls. (n for WT sed=3, WT run=6, HET sed=4, HET run=3, DES sed=4, DES run=4). Group differences were assessed via a two-way ANOVA, error bars indicate ± SD.

### Proteomics

Fractional synthesis of individual proteins was measured via LC-MS and clustered through the NIH DAVID database. Because we did not see relevant differences between WT and HET animals in histopathology and the protein levels we investigated, we only analyzed proteomics in WT and DES samples. Data filtering and calculations were performed according to previous reports^28^. Baseline protein synthesis across all proteins tended to be lower in WT than in DES in the control condition (0.151 ± 0.174 to 0.168 ± 0.167) but did not differ significantly (p=0.97). However, with running global protein synthesis increased to a greater extent in WT (on average 36%) (p<0.05) compared to DES (on average 20%) (p=0.19). Running increased the synthesis of 32 out of 45 proteins in WT and decreased protein synthesis in 8 out of 45 proteins, with 5 remaining unchanged. In DES an equal number of proteins increased their synthesis with running (32 out of 45). However, out of the 32 proteins that increased in WT with running only 26 also increased in DES. Of the proteins that showed the strongest difference in synthesis between WT and DES, many were associated with mitochondrial function (l-lactate dehydrogenase A chain (Ldha), 3-mercaptopyruvate sulfurtransferase (Mpst)), glycolysis (glucose-6-phosphate isomerase (Gpi), phosphoglycerate kinase 1 (Pgk1)) and muscle contraction (myosin light chain 1/3, skeletal muscle isoform (Myl1), myosin light chain 3 (Myl3), myosin-4 (Myh4)). Interestingly, among those proteins that showed a strong increase in DES but not in WT were proteins that are closely associated with muscle contraction such as Tropomyosin beta chain (Tpm2), Troponin T, fast skeletal muscle (Tnnt3) and Tropomyosin alpha-1 chain (Tpm1).

Out of 22 potential annotation clusters, mitochondrial- and glycolytic protein clusters were among the most prominent in our samples (Fig5A and B). We found 17 mitochondrial proteins and 12 glycolytic proteins that passed our quality control criteria (selected via GOTERM_CC_DIRECT and KEGG_PATHWAY, respectively). Four out of 17 mitochondrial protein synthesis values were significantly altered in WT through running, while no statistical difference could be found for DES animals (Figure 5A). The protein showing the strongest increase with running in the WT group was cytochrome c, somatic (Cycs) (4.3-fold as high) (p<0.001), while peroxiredoxin 1 (Prdx1) was the protein with the strongest albeit insignificant decrease in protein synthesis (27%). For DES, ATP synthase, H+ transporting, mitochondrial F1 complex, beta polypeptide (Atp5b) was the protein that showed the strongest tendency for an increase with running without reaching statistical significance (p=0.14). Another mitochondrial protein that stood out was heat shock protein family D member 1 (Hspd1), which decreased by 11% with running in WT animals but by 67% in DES animals compared to their own control condition as well as by 88% compared to the WT control condition.

**Figure 5 –.**
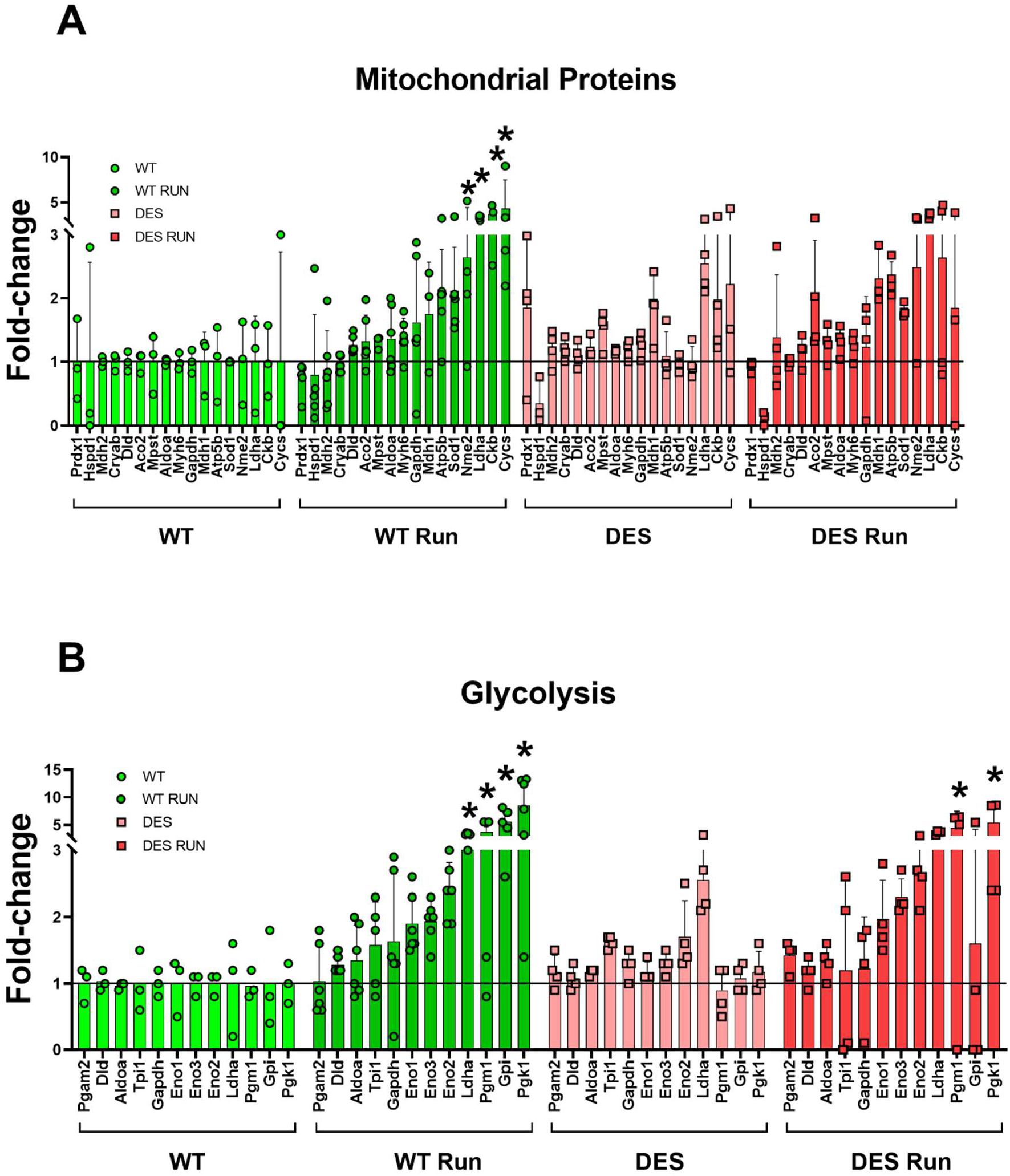
Change in mitochondrial and glycolytic protein synthesis with chronic exercise. Mitochondrial (Figure 5A) and glycolytic proteins (Figure 5B) were clustered according to the NIH database DAVID (using GOTERM_CC_DIRECT and KEGG_PATHWAY selections). All fractional synthesis values of proteins were normalized to WT control. Four out of 17 mitochondrial protein synthesis values were significantly altered through running in WT. No mitochondrial protein synthesis values were significantly altered in DES (Figure 5A). Four out of 12 glycolytic protein synthesis values were significantly altered through running in WT. Two glycolytic protein synthesis values were significantly altered in DES (Figure 5B). (n for WT sed=3, WT run=6, DES sed=4, DES run=4)* indicates a p-value of >0.05 compared to their own sedentary control (post hoc analysis). Error bars indicate ± SD

For the synthesis of glycolysis associated proteins, four out of 12 changed significantly (post-hoc analysis) with running in WT while only two changed in DES (Fig5B). The most robust changes with running occurred in Pgk1 (8.6-fold as high) (p<0.0001), Gpi (5.6-fold as high) (p<0.0001) and Pgm1 (3.8-fold as high) (p<0.05) in the WT animals. In the DES animals, the proteins showing the strongest increase with running were Pgk1 (p<0.001) and Pgm1 (p<0.01) when compared to their own control condition.

Finally, looking at proteins directly involved in muscle contraction we found substantial differences (Figure 6). Particularly WT and DES showed different adaptive responses to the running protocol: Myl3 protein synthesis increased to a level 2-fold as high in running WT compared to sedentary WT (Fig6A). In contrast, in DES animals running decreased Myl3 by 7%. This discrepancy can be partially attributed to the fact that baseline Myl3 levels were 2.2-fold higher in DES compared to WT. Following a similar pattern, Myh4 increased by 23% with running in the WT group but decreased by 16% in the DES group (Fig6B). For myosin-6 (Myh6), running increased protein synthesis by 39% in WT and only by 6% in DES (Fig6C). Similarly, myosin-7 (Myh7) protein synthesis increased by 39% with running in WT and by 7% in DES (Fig6D).

**Figure 6 -.**
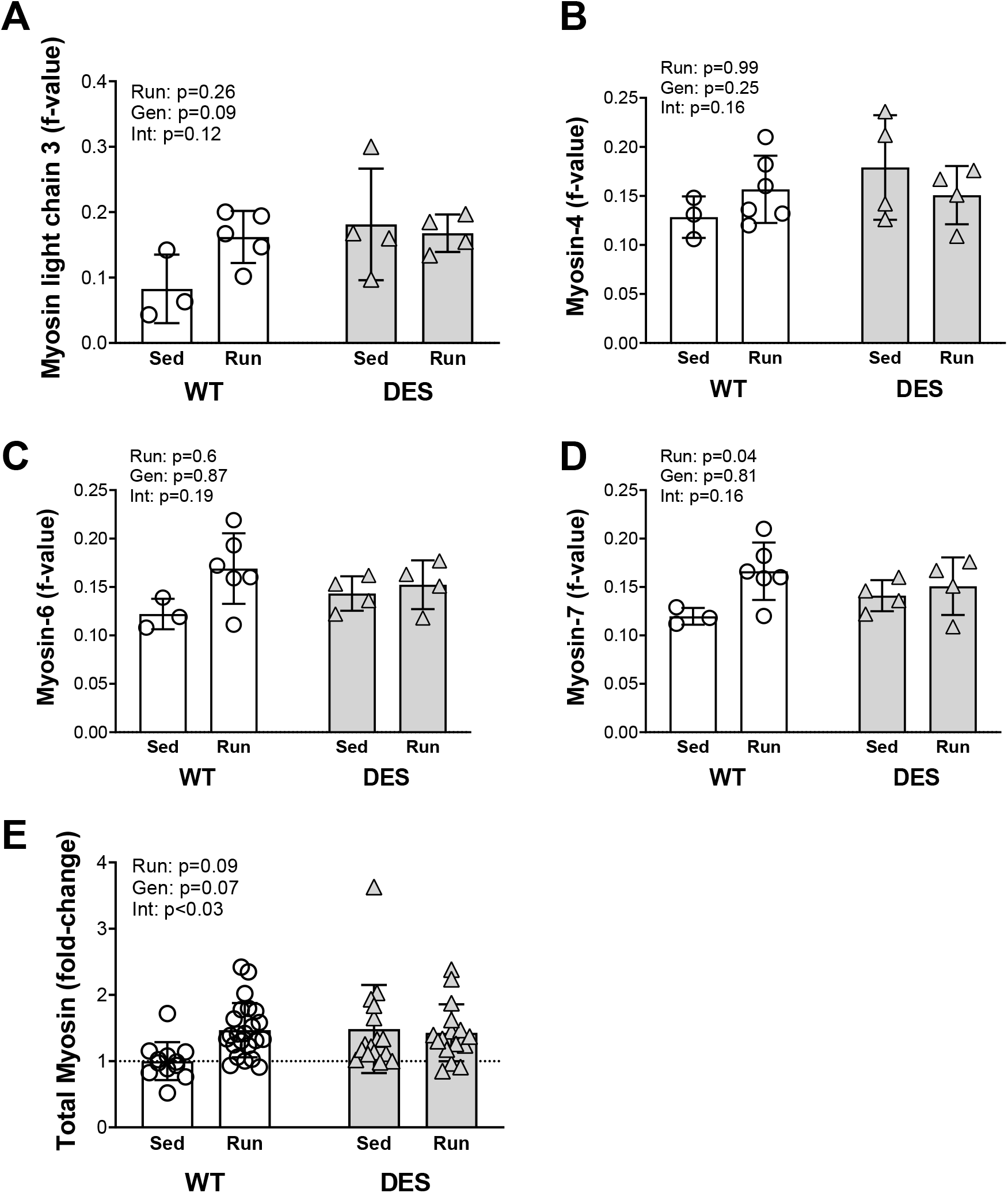
Fractional synthesis of different myosin isoforms in the gastrocnemius muscle with chronic exercise. Displayed are raw f-values of the individual proteins: myosin light chain 3 (Myl3) (A), myosin-4 (Myh4) (B), myosin-6 (Myh6) (C) and myosin-7 (Myh7) (D). Values of all myosin isoforms analyzed in (A-D) were normalized to the WT control condition and are displayed as fold-change as a clustered analysis of myosin isoforms (E). (n for WT sed=3, WT run=6, DES sed=4, DES run=4). Group differences were assessed via a two-way ANOVA and p-values are listed on the graphs. Error bars indicate ± SD Gen=Genotype, Run=Running, Int=Interaction; Sed=sedentary controls.

Based on the distinct pattern in protein synthesis between WT and DES, we grouped the myosin isoforms from Figure 6 A-D together and normalized all values to the WT control group (Fig6E). This overview revealed a significant interaction effect between the genotype and running (p<0.03): Clustered myosin protein synthesis values increased by 47% with running in the WT group. In DES, however, protein synthesis at baseline were already 49% higher than in WT control animals and slightly decreased (6%) from there through running.

## DISCUSSION

We recently created a novel CRISPR-Cas9 rat model for the most common type of desminopathy^26^. While the age of onset in the human disease ranges from infancy to late adulthood, most patients first develop symptoms after their third decade^4–6,15^. In previous studies we investigated histopathological and molecular changes in our model at a preclinical age after functional overload and in old rats after acute eccentric loading. We showed that mutant animals at an age corresponding to a preclinical age in humans had a mild myopathy and increased markers for muscle injury and repair^26^. To see whether ageing exacerbates the phenotype, in our next study we looked at 400–500-day old animals in conjunction with exercise. Interestingly, while signs of chronic remodeling appeared similarly pronounced in old, compared to young, desmin mutants, DES animals appeared to be protected from exercise induced muscle damage unlike their WT littermates^27^. We concluded that ageing alone does not manifestly aggravate the disease progression, and hypothesized that our rats, having spent their lives in cages in a mostly sedentary manner, lack an adequate stimulus to trigger the pathophysiology. In this study, we set out to examine the effects of chronic injury on DES muscle. We chose downhill treadmill running to provide repeated mechanic stress and stimulate eccentric-induced injury and regeneration in the muscles of the hindlimbs while minimizing the metabolic cost of the exercise.

During downhill running, plantar flexor muscles (gastrocnemius muscle, soleus muscle and plantaris muscle) primarily perform eccentric contractions to decelerate the animal’s center of mass in order to maintain a constant running velocity^29^. These eccentric contractions cause injury consisting of disruptions of sarcoplasmic reticulum and other structures followed by local inflammatory and regenerative processes in the activated muscles^29–31^. At the same time, running down an incline leads to lower whole body oxygen consumption, blood lactate concentrations and presumably cardiovascular strain than during level locomotion at the same speed^32–34^.

From previous reports on downhill treadmill exercise that had 400-to 500-g wildtype Sprague-Dawley rats run continuously at 16 m/min for 90 min either on the level or down a 16° incline^29^ we expected much longer running times in our animals. The longest time our rats ran was 22 min at a top velocity of 17m/min, while the average maximal running time in our WT was 17.5 min at a median top velocity of 13 m/min. Our animals were heavier, at an average body weight of 618g, but we saw no correlation between body weight and running time (R squared = 0.25; data not shown). Even though the study by Armstrong et al. did not report the age of their animals, given our rats were substantially heavier it appears likely that our animals may have been older. Other reports have shown a decrease in running endurance with increasing age as well as a pronounced stress response and poorer compliance in treadmill running in older animals compared to younger rats^35^. The difference between previously reported running performance and the running endurance of our rats might therefore be age-related. However, we did not see signs of an increased stress response as monitored through weekly body weight, and compliance was good after two weeks of familiarization.

One DES animal had to be terminated during the course of the running experiments in week 2 due to severe bowel obstruction. Bowel obstruction or intestinal pseudo-obstruction have been reported in rare human cases of desminopathy, and histopathologic assessment of the affected intestines showed subsarcolemmal aggregates typical for the disease in smooth muscle^36^.

We saw distinct differences between the genotypes in their running performance. Performance in DES rats tended to be lower at the beginning of protocol, the running endurance in DES rats dropped significantly by the fourth ILT, and performance continued to decrease for the whole 24-day training period. Given this stark decrease in muscle function, it was somewhat surprising to see only moderate histopathological changes after running in the DES animals. The R349P animals demonstrated a higher percentage of fibers with central nuclei in DES compared to WT rats in the sedentary condition, which is in line with our previous findings in this strain^26^. However, this percentage did not go up in DES animals after the downhill running protocol, but rather tended to be reduced. In our previous study on acute bouts of eccentric loading in R349P rats, we also did not see a significant increase in fibers with central nuclei both in WT and DES, which we attributed to the time course of our experiment, as the centralization of myonuclei is thought to be a sign of regeneration following ~7 days after injury^37^ and we collected the tissue 24 hours after loading. This explanation obviously does not hold up in the current chronic injury study. To assess myofibrillar injury via membrane damage, we stained for intracellular IgG. Except for one DES animal with 0.13% IgG-positive fibers, no animals showed any signs of membrane permeability at the time of collection. In our previous studies, we saw no IgG-positive fibers at baseline, but an increase in IgG-positive fibers after eccentric loading that was 5-fold higher in older WT than in age-matched DES animals.

Molecular markers for injury and repair followed a similar pattern as we previously saw in preclinical animals and after acute eccentric loading. Most notably, mLIM was upregulated in sedentary HET and more so in DES animals compared to WT, and while mLIM levels were increased in WT and HET after running, they did not change significantly in DES. mLIM has been found to interact with many different proteins both in the cytoplasm and the nucleus, and therefore serves a variety of functions. One of its integral roles is at the Z-disk, where it has been suggested to act as a scaffold protein that promotes the assembly of macromolecular complexes along sarcomeres and the actin-based cytoskeleton as well as a mediator of stretch signaling^38^. An upregulation of this Z-band associated protein after eccentric stress would be in line with previous findings that eccentric exercise has been found to lead to muscular adaptation specifically around the Z-band^39^. Dysferlin, by contrast, is a sarcolemmal protein that is mainly involved in membrane repair by facilitating the fusion of repair vesicles with a damaged membrane^40–42^. Therefore, it is usually upregulated after membrane injury. We saw a tendency for Dysferlin to be higher in DES animals compared to WT and this difference we accentuated after 6 weeks of running. As we do not see any indication for membrane injury as assessed via IgG-staining, we assume that this increase is due to an accumulation of Dysferlin in protein aggregates. Subsarcolemmal protein aggregates are a main histopathological hallmark in Desminopathy, and while the aggregates are initiated through the accumulation of misfolded desmin, they also contain debris of other proteins, including dysferlin^23^. We saw a similar trend towards higher desmin levels, both histologically as an increase in desmin aggregates and biochemically as an increase in western blots after running in the DES animals. Overall, we conclude that disease progression in DES rats was aggravated by downhill running, but not, as we expected, through mechanical stress-induced membrane damage.

There is conflicting data on the effects of mechanical stress on desminopathic muscle. While most reports point towards an increased mechanical vulnerability of mutated myofibers^43,44^, there is also evidence that loss of desmin might render muscle fibers less susceptible to membrane damage^45–47^. However, to our knowledge there are no studies investigating the influence of chronic mechanical stress in desminopathy. There is anecdotal evidence that desminopathic patients who exercise suffer from a worse disease progression^48^, but this is difficult to interpret since the same mutation in desmin can present varied severity and progression.

This led some investigators to hypothesize that stiffness and mechanical stress is probably not the most important kind of stress that leads to disease progression in desminopathy^48^. The fact that creatine kinase (CK) serum levels, a classic, albeit crude, marker of muscle damage^49–51^, is of limited diagnostic value might be another indicator that membrane damage is not prominent in desminopathy. Fewer than 60 % of mutation carriers have elevated CK levels, and a third of patients with manifest skeletal muscle disease were reported to have normal CK levels^52^.

It is important to consider the fact that given their earlier exhaustion, the DES animals ran a significantly shorter distance than WT and HET over the 6-week training period and therefore received a lower absolute stimulus. While WT rats ran for an average of 171 m per ILT, HET rats ran 133 m and DES only 82 m. Still, this stimulus was enough to almost double the amount of desmin aggregates.

Desmin-positive aggregates are thought to be a central disease-driving mechanism. They are caused by misfolded desmin that cannot be degraded by the PQC. Over time, other cellular debris gets entangled in the aggregates that are then thought to “clog” autophagic and regenerative mechanisms. We therefore investigated proteins involved in autophagy.

ULK1 plays a pivotal role in the initiation of autophagy. Its activity is positively regulated through phosphorylation at Ser^555^ and negatively regulated through phosphorylation at Ser^757 53–55^. Single bouts of exercise have been shown to increase ULK1 phosphorylation at Ser^555^ both in mice and humans^56,57^, but phosphorylation both at Ser^555^ and Ser^757^ did not change after a long-term resistance exercise protocol in rats^58^. We saw lower phosphorylation of ULK1 both at Ser^555^ and Ser^757^ in DES animals compared to WT, and phosphorylation went down in all genotypes after running. It is not clear whether this is a long-term adaptive process, possibly to reduce autophagy to sustain anabolism and muscle growth in a response to training, or a short-term effect due to collection time. Autophagy responses after exercise peak at about 2 h post exercise and dwindle after 3–4 h post exercise^59^. We collected muscles 24 h after their last running session making it unlikely that ULK1 levels reflected the last exercise bout. Downstream from ULK1, we also did not identify a stark impairment in autophagy in DES animals. However, total ubiquitination of proteins as a marker of ubiquitin proteasome-system (UPS) activity, went down in WT and up in DES animals, which might indicate impaired protein degradation or an overactive UPS response.

Our previous investigation of R349P mutant rats showed histological and molecular signs of increased muscle remodeling at baseline compared to WT. However, acute exercise did not seem to exacerbate this effect and DES muscle turned out to be less vulnerable to injury than WT muscle^26^. In the study at hand, we exposed rats to a chronic running stimulus and combined this with stable isotope labeling to investigate whether repeated exercise can 1) challenge the protection of R349P muscle to injury, 2) elicit a distinct, adaptive response in WT compared to DES animals, and 3) determine which proteins are synthesized in WT and DES animals.

In line with our previous study, we found that global protein synthesis values in the sedentary condition were slightly higher in DES compared to WT rats. The chronic exercise stimulus elevated global protein synthesis values in the WT group to a greater level than in the DES group. Consequently, the fact that most protein synthesis values showed a substantially higher fold-change with exercise in WT can be explained two ways: lower baseline levels and a greater adaptive response to running in WT. Part of the latter could be interpreted as a function of the aforementioned greater absolute training stimulus in WT. However, in healthy individuals relative exercise intensity and volume are likely more relevant for eliciting molecular responses and adaptations to exercise than absolute ones. This becomes evident when looking at the fact that beginners commonly show amplified molecular signaling after exercise compared to more experienced athletes, despite substantially lower training intensities^60^. Since the relative intensity and the running volume for the mutant animals was identical to their WT littermates, a lack of stimulus appears unlikely to explain the differences between the genotypes. In addition, many of the protein synthesis values in DES at baseline were close to or at the same level as in WT with chronic exercise. That DES did not show a further increase with exercise could therefore be a sign of a physiological ceiling of protein synthesis rates rather than a lack of adaptation.

Investigating proteins individually or as part of a cluster revealed that DES mutant animals tended to synthesize different proteins than WT controls both at baseline and following chronic exercise. For example, running increased cytochrome c 4.3-fold in WT, whereas in DES mutants the same protein was elevated in the control condition (2.2-fold compared to WT control) and stayed virtually unchanged with exercise (1.9-fold). In contrast, peroxiredoxin 1 was synthesized almost 2-fold more at baseline in DES compared to WT, but decreased with running to a level equivalent to WT. Peroxiredoxin is thought to be involved in the protection of cells from oxidative stress and is elevated in inflammatory scenarios and cancer^61,62^. Higher synthesis values in DES at baseline could therefore indicate a greater need for coping with reactive oxygen species induced by the mutation, that is partially relieved through exercise.

For glycolytic proteins, synthesis values at baseline were more comparable between WT and DES than mitochondrial proteins. However, in response to the chronic exercise stimulus we could still see distinct responses between the two genotypes. One of the most pronounced discrepancies was the synthesis of glucose-6-phosphate isomerase. While baseline values were similar between WT and DES, running elevated synthesis to levels 5.6-fold in WT while in DES synthesis of the same protein rose only to levels 1.6-fold as high (largely due to a single individual). In our acute study of R349P mutants after exercise, we found a diminished glucose tolerance in DES animals (accepted for publication). Furthermore, in an investigation of muscle metabolism after nerve damage we found intramuscular glucose to be elevated despite a tendency for lower glucose 6 phosphate levels^63^. Similar shifts in the levels of muscle metabolites have been observed for other congenital muscular dystrophies such as dysferlinopathy and Duchenne muscular dystrophy^64–66^. The disturbance of systemic as well as local muscle glucose metabolism in our desminopathy model and other muscular dystrophies/atrophies could point to a common physiological mechanism by which skeletal muscle copes with challenges to its integrity, despite different genetic and environmental reasons underlying such challenges.

Finally, looking at contractile protein isoforms individually and in a clustered manner we found baseline protein turnover to be elevated in DES compared to WT. This is in line with histological observations in this and our earlier papers, where we found an increased number of central nuclei in DES muscle at baseline. The finding of a higher number of central nuclei in combination and slightly elevated protein synthesis via puromycin, and a decreased susceptibility to exercise induced muscle damage in an acute setting led us to hypothesize that the R349P mutation in desmin might have caused an adaption in DES animals that results in higher contractile protein turnover. Indeed, in this study we found that DES animals have almost 50% higher values of contractile protein turnover at baseline compared to WT animals. While chronic exercise elevated contractile protein turnover in WT (~50%), in DES animals the running did not significantly alter contractile protein turnover.

In summary, in this study we challenged desminopathic muscle with a chronic, physiological exercise stimulus. We found a decreased running capacity in animals affected by the mutation, which was further exacerbated over the course of the study. In accordance with previous studies, we found histological signs of increased fiber damage and remodeling as well as a shift in fiber distribution towards smaller fibers in desminopathy. We found desmin aggregates at baseline in the mutant rats which become more pronounced with exercise training. On a protein level we found indicators of increased autophagic and proteasome activity with running in DES compared to WT. Finally, stable isotope labeling and LC-MS analysis revealed distinct adaptations of the proteomes of WT and DES animals at baseline as well as with exercise. Most prominently, key proteins of mitochondria, glycolysis and thick filaments increased their synthetic activity robustly with running in WT, while DES showed higher baseline values and little change in response to running.

Future studies will have to determine whether there is a healthy dose of exercise for R349P mutant muscle and its human analog, or whether physical stress is to be avoided under any circumstance. Moreover, the mechanical association of desmin with other force transfer proteins must be further elucidated to find potential targets to ameliorate the intrinsic stress desmin-mutant muscle appears to suffer from.

## METHODS

### CRISPR-Cas9-mediated knock-in

Using CRISPR-Cas9, the missense mutation *DES* c.1045-1046 (AGG > CCG) was introduced in exon 6, leading to p.R349P in rats. For a detailed description of the generation and characterization of this rat desminopathy model see Langer et al, 2020^26^. Male rats between 400 to 560 days of age were selected to study functional, histological, and biochemical differences in the phenotype of CRISPR-Cas9 created knock-in animals carrying a mutation in desmin (DES; n = 9), heterozygous (HET; n = 8) or healthy, wild type littermates (WT; n = 9). Animals were housed in 12:12 h light–dark cycles and fed ad libitum.

### Downhill running

The downhill running protocol was performed on a motorized treadmill (Exer-3/6, Columbus Instruments, Columbus, OH) with an adjustable belt speed (3–100 m/min). The speed of the treadmill was calibrated at the beginning of each week of the experiment. The rear of the treadmill was equipped with a low-voltage, electric stimulating grid with individual on/off switches per lane. The stimulating grid was set to deliver 0.2 mA at a frequency of 1 Hz.

### Familiarization with treadmill and run to exhaustion

Rats from each genotype were randomly assigned to a “sedentary” (SED) or an “exercise” (RUN) group. None of the rats had been on a treadmill or subjected to any kind of exercise prior to familiarization. Rats in the exercise group (n=14) were accustomed to treadmill running over two weeks, starting at 10m/min at a 0° decline. To encourage running, rats were prodded with a soft bristled brush when they fell back onto the back quarter of the treadmill. The exercise session lasted until exhaustion, which was defined as the rat’s inability to maintain running speed despite repeated physical prodding and staying on the electrical grid for more than five consecutive seconds^67^. At exhaustion, the electrical grid was immediately switched off and the animal removed from the treadmill. This protocol was performed every other day for eight days. On day nine, the protocol was set to 15m/min at a 0° decline, followed by a day of rest. On day eleven of familiarization, speed was set to 5 m/min and increased by 2 m/min every 3 min until exhaustion. This was followed by three days of rest, after which the downhill running protocol was started (Figure 1A).

### Downhill running protocol

Rats in the exercise group were subjected to downhill running at a decline of −10° for four weeks. For three weeks, rats performed an incremental load test (ILT) on day 1, 3 and 5, starting at a running speed of 5 m/min, which was increased by 2 m/min every 3 min until exhaustion. Upon exhaustion, the maximal speed completed as well as total running time were recorded. On day 2 and 4, to avoid overexercising but maintain a state of chronic physical activity, rats were exposed to walking at 40% of their respective maximal completed running speed for 10 minutes. On day 6 and 7, rats were rested. Weight was recorded on day 1 of each week. In week 4, ILT was performed on days 1 and 3, 40%-speed running was performed on days 2 and 4. On day 5, rats ran at 15m/min on 0° decline to exhaustion and total running time was recorded. Animals were euthanized 24h after the final running test.

### Deuterium oxide labelling

Animals were stable isotope labeled with deuterium oxide using previously published protocols with modifications^68,69^. Briefly, endogenous deuterium levels were raised to about 4 % by a priming intraperitoneal injection of 0.014 mL *g−1 bodyweight deuterium oxide (99.8 % + Atom D, Euriso-Top GmbH Saarbrücken) and 0.9 % NaCl. Deuterium levels were maintained via drinking water (4 % deuterium oxide). The stable isotope labeling started five days before collection and an independent group of unlabeled rats served as a control.

### Muscle collection

Rats were anaesthetized via isoflurane inhalation (2.5%). The gastrocnemius (GAS), soleus (SOL) and tibialis anterior (TA) muscles were excised from the left hindlimb, blotted dry, and weighed. Subsequently, the muscles were pinned on cork at resting length and frozen in liquid nitrogen-cooled isopentane for histological and biochemical analyses. Liver, heart and serum were also collected. On completion of tissue removal, rats were euthanized via cardiac puncture.

### Histology

Frozen muscles were blocked, and serial cross-sections (10 μm) were cut using a Leica CM 3050S cryostat (Leica Microsystems, Buffalo Grove, IL, USA). For hematoxylin and eosin (H&E) staining, sections were brought to room temperature, then consecutively submerged in EtOH 90% and tap water for 30 s. Slides were then incubated in Mayer’s hemalum solution (Merck, Darmstadt, Germany) for 6 min, blued under running lukewarm tap water for 2 min, and then dipped in tap water for another 8 min. Slides were incubated in 0.5% aqueous eosin γ-solution (Merck, Darmstadt, Germany) for 3 min, then dehydrated using 90%, 90%, and 100% EtOH for 1 min each. Alcohol was removed by submerging sections in Xylene for 5 min. After sections were air dried, they were mounted with DPX Mountant for histology (Sigma, Darmstadt, Germany).

For immunohistochemistry, muscle sections were fixed in acetone for 5 min at −20°C, followed by 3 × 5-minute phosphate-buffered saline washes in 0.1% Tween-20 (PBST) before blocking with 5% natural goat serum (NGS) for 30 min at room temperature. Sections were then incubated with primary antibody overnight at 4°C. For detection of desmin aggregates, a monoclonal desmin antibody (mouse, IgG2a, Santa Cruz Biotechnology (Dallas, Texas), 1:200 in NGS) was used. The sarcolemma was stained using a polyclonal laminin antibody (rabbit, IgG (H+L), Sigma Aldrich (St Louis, Missouri), 1:500 in NGS). After incubation in primary antibody, sections were washed three times with phosphate-buffered saline containing 0.1% Tween-20 for 5 min and then incubated in secondary antibody (goat anti-rabbit AlexaFluor® 647, Thermo Fisher Scientific (Waltham, Massachusetts), Lot 1910774 for laminin, goat anti-mouse Alexa Fluor® 488, Thermo Fisher Scientific, Lot 1900246 for desmin) for 30 min at room temperature. Sections were mounted using ProLong Gold Antifade reagent (Life Technologies). Membrane damage was assessed using anti-IgG staining using an anti-rat antibody (IgG H+L AlexaFluor® 647 1:100, Thermo Fisher Scientific, Lot 2005938 in PBST for 2h at room temperature). Slides were imaged using a Leica DMi8 inverted microscope using the HC PL FLUOTAR 10x/0.32 PH1 objective (Leica Microsystems, Wetzlar, Germany) and using the LAS X software. For comparative analysis, exposure length remained fixed for all samples.

Muscle fiber properties and fiber types were analyzed using FIJI and SMASH (MATLAB) software from the whole muscle sections.

### Hydroxyproline determination of collagen content

Collagen content was determined using a hydroxyproline assay^70^. Muscle tissue was placed in 1.7ml tubes and weighed for wet mass. Tissue was dried on a heating block with lids open for 30 min at 120°C. Each sample was then weighed for dry mass and hydrolyzed in 200 µL of 6N HCl at 120°C for 2 hours before drying for 1.5 hours at 120°C. The dried pellet was resuspended in 200 µL of hydroxyproline buffer (173 mM citric acid, 140 mM acetic acid, 588 mM sodium acetate, 570 mM sodium hydroxide). The sample was further diluted 1:67 in hydroxyproline buffer. 150 µL of 14.1 mg/ml Chloramine T solution was added to each sample, then vortexed and incubated at room temperature for 20 minutes. 150 µL aldehyde-perchloric acid containing 60% 1-propanol, 5.8% perchloric acid, and 1M 4-(dimethylamino)benzaldehyde was then added to each sample, before vortexing and incubating at 60°C for 15 minutes. The tubes were then cooled for 5-10 minutes at room temperature. Samples were read at 550 nm on an Epoch Microplate Spectrophotometer (BioTek Instruments Limited, Winooski, VT). Hydroxyproline was converted to collagen mass assuming hydroxyproline contributes to 13.7% of the dry mass of collagen^71^.

### Immunoblotting

Muscles were powdered on dry ice. Frozen muscle powder was homogenized in 200 μL sucrose lysis buffer (50 mM Tris pH 7.5, 250 mM sucrose, 1 mM EDTA, 1 mM EGTA, 1% Triton X-100, 1% protease inhibitor complex) on a vortexer for 60 min at 4°C. Following centrifugation at 10 000 g for 10 min, the supernatant was collected. Protein concentrations were determined in triplicates using the DC protein assay (Bio-Rad, Hercules, CA, USA) and sample concentrations were adjusted to 1 μg protein/μL. Protein aliquots were denatured in Laemmli sample buffer at 100°C for 5 min and then loaded (10– 15 μg protein per lane) on 4–20% Criterion TGX Stain-free gels (Bio-Rad) and run for 45 min at 200 V. Total protein was visualized after a 1 min exposure to UV light that induced a fluorescent reaction from the proteins in the gel. Following quantification of total protein, proteins were transferred to PDVF membrane at 100 V for 30–60 min, depending on the size of the protein of interest. Efficient transfer was confirmed using Ponceau staining of the membrane. Membranes were then washed and blocked in 1% fish skin gelatin dissolved in Tris-buffered saline with 0.1% Tween-20 for 1 h. Membranes were then rinsed and probed with primary antibody overnight at 4°C. The next day, membranes were washed and incubated with HRP-conjugated secondary antibodies at 1:7500 to 1:10 000 in 1% skim milk-Tris-buffered saline with 0.1% Tween-20 for 1 h at room temperature. Immobilon Western Chemiluminescent HRP substrate (Millipore, Hayward, CA, USA) was then applied to the membranes for protein visualization by chemiluminescence. Image acquisition and band quantification were performed using a ChemiDoc MP System with Image Lab 5.0 software (Bio-Rad). Protein levels of each sample were calculated as band intensities relative to total protein. The following antibodies were used in this study at a concentration of 1:1000: *Cell Signaling* (Cell Signaling Technology, Danvers, MA: ribosomal protein S6 (Ser240/244) (# 5364), microtubule-associated proteins 1A/1B light chain 3B (LC3B) (#2775; lot 10), heat shock protein beta-1 (HSP27) (#2402; lot 8), heat shock protein HSP 90-alpha (HSP90) (#4877; lot 5), unc-51 like autophagy activating kinase 1 (ULK1) (Ser757) (#14202; lot 1); p62 (#5114; lot 4); Annexin A2 (#8235; lot 2), caspase 3 (#9662), autophagy related 7 (Atg 7) (#8558); NOS (pan) (#2977); *Santa Cruz* (Santa Cruz Biotechnology Inc., Dallas, TX): Dystrophin (#365954; lot E2711), Dysferlin (#16635; lot H162), Desmin (#271677; lot F1913), muscle LIM protein (mLIM)/cysteine and glycine-rich protein 3 (mLIM) (#166930; lot E2814); septin 1 (#373925); *Millipore Sigma* (Merck Group): insulin receptor substrate 1 (IRS1) (#06-248; lot 2465193); *Abcam* (Abcam Inc., Eugene, OR): total oxidative phosphorylation (OXPHOS) (MS604-300).

### Proteomics

#### Body water enrichment analysis

Rat serum and liver were distilled overnight upside down on a bead bath at 85°C to evaporate out body water. Deuterium present in the body water were exchanged onto acetone, and deuterium enrichment in the body water was measured via gas chromatography mass spectrometry (GC-MS)^72^.

#### Tissue preparation for LC-MS

Tissues were homogenized in homogenization buffer (100 mM PMSF, 500 mM EDTA, EDTA-free Protease Inhibitor Cocktail (Roche, catalog number 11836170001), PBS) using a 5 mm stainless steel bead at 30 hertz for 45 seconds in a TissueLyser II (Qiagen). Samples were then centrifuged at 10,000 g for 10 minutes at 4°C. The supernatant was saved and protein was quantified using a Pierce BCA protein assay kit (ThermoFisher, catalog number 23225). 100 ug of protein was used per sample. 25 µL of 100 mM ammonium bicarbonate solution, 25 µL TFE, and 2.3 µL of 200 mM DTT were added to each sample and incubated at 60°C for 1 hour. 10 µL 200 mM iodoacetamide was then added to each sample and allowed to incubate at room temperature in the dark for 1 hour. 2 µL of 200 mM DTT was added and samples were incubated for 20 minutes in the dark. Each sample was then diluted with 300 µL H2O and 100 µL 100 mM ammonium bicarbonate solution. Trypsin was added at a ratio of 1:50 trypsin to protein (trypsin from porcine pancreas, Sigma Aldrich, catalog number T6567). Samples were incubated at 37°C overnight. The next day, 2 µL of formic acid was added. Samples were centrifuged at 10,000 g for 10 minutes, collecting the supernatant. Supernatant was dried by speedvac and re-suspended in 50 µL of 0.1 % formic acid/3% acetonitrile/96.9% LC-MS grade water and transferred to LC-MS vials to be analyzed via LC-MS.

#### LC-MS analysis

Trypsin-digested peptides were analyzed on a 6550 quadropole time of flight (Q-ToF) mass spectrometer equipped with Chip Cube nano ESI source (Agilent Technologies). High performance liquid chromatography (HPLC) separated the peptides using capillary and nano binary flow. Mobile phases were 95% acetonitrile/0.1% formic acid in LC-MS grade water. Peptides were eluted at 350 nl/minute flow rate with an 18-minute LC gradient. Each sample was analyzed once for protein/peptide identification in data-dependent MS/MS mode and once for peptide isotope analysis in MS mode. Acquired MS/MS spectra were extracted and searched using Spectrum Mill Proteomics Workbench software (Agilent Technologies) and a mouse protein database (www.uniprot.org). Search results were validated with a global false discovery rate of 1%. A filtered list of peptides was collapsed into a nonredundant peptide formula database containing peptide elemental composition, mass, and retention time. This was used to extract mass isotope abundances (M0-M3) of each peptide from MS-only acquisition files with Mass Hunter Qualitative Analysis software (Agilent Technologies). Mass isotopomer distribution analysis (MIDA) was used to calculate peptide elemental composition and curve-fit parameters for predicting peptide isotope enrichment based on precursor body water enrichment (p) and the number (n) of amino acid C-H positions per peptide actively incorporating hydrogen (H) and deuterium (D) from body water. Details of fractional synthesis calculations and data filtering criteria have been described in detail in Holmes et al., 2015^28^. Proteins were clustered with the Functional Annotation Tool of the NIAID/NIH Database for Annotation, Visualization and Integrated Discovery (DAVID) version 6.8^73^.

#### Statistics

Appropriate sample sizes to detect group effects were estimated on the basis of previous experiments with the same strain of animals^26,27^. Two-way analysis of variance (ANOVA) with a post hoc Tukey’s multiple comparisons test was used to test the null hypothesis. An alpha of *P* < 0.05 was deemed statistically significant, and a *P* value between 0.05 and 0.1 was called a trend. Biological replicates are reported as “n” in each figure legend. “Genotype” = “Gen” indicates effects of wildtype (WT), heterozygous (HET) and desmin R349P mutant (DES) genotypes on the test. “Running” = “Run” indicates effects between sedentary (SED) or exercised (RUN) group. “Interaction” = “Int” indicates the interaction effect between “genotype” and “running”. Desmin aggregates (% of whole section) in Figure 2F were tested via unpaired t-test. Effect sizes for proteomics were calculated according to Hedges’ g^74^. Data in the text are reported as mean ± standard deviation, and data in the figures are visually represented as scatter dot plot with error bars indicating standard deviation. All analysis was performed with GraphPad Prism Version 8 (La Jolla, CA, USA).

#### Animals and ethical approval

All procedures were approved by the Institutional Animal Care and Use Committee of the University of California, Davis, which is an AAALAC-accredited institution. Animal housing was in accordance with recommendations of the Guide for the Care and Use of Laboratory Animals.

## Author contributions

AAM and HTL conceptualized and wrote the paper. AAM and HTL take a shared first authorship as they conceptualized the project together and conducted the bulk of the experiments (AAM: treadmill experiments, histology and immunoblotting, HTL: immunoblotting and dynamic proteomics data analysis), and as AAM conducted the primary animal experiments, she is listed first. AB processed tissue samples for the LC-MS and contributed to the data analysis of dynamic proteomics. AMA conducted the hydroxyproline experiments. HZF and MH contributed to conceptualization of the study and revision of the manuscript. KB supervised the project and contributed to conceptualization of the experiments, data analysis and revision of the manuscript.

## Acknowledgements

The research was made possible by a generous gift from the Bertin-Barbe Family. The work was further supported by the National Institute of Aging through project grants (R01AG056999 and R01AG45375). Agata Mossakowski was supported by a postdoctoral fellowship from Deutsche Forschungsgemeinschaft. We would like to thank Diana L. Young, Nichole L. Anchell, Kayla M. Jager, Phuong T. Dao and the other technical staff at the UC Davis Mouse Biology Program for their work in generating this rat model. We would like to thank Dr. Lucas Smith for his SMASH software and use of his microscope.

Agata Mossakowski, Henning Langer, Alec Bizieff, Alec M. Avey, Hermann Zbinden-Foncea, Marc Hellerstein and Keith Baar all declare that they have no conflict of interest.

## REFERENCES

1. Bär H, Strelkov S V., Sjöberg G, Aebi U, Herrmann H. The biology of desmin filaments: How do mutations affect their structure, assembly, and organisation? J Struct Biol. 2004. doi:10.1016/j.jsb.2004.04.003

2. Clemen CS, Herrmann H, Strelkov S V, Schröder R. Desminopathies: pathology and mechanisms. Acta Neuropathol. 2013;125(1):47–75. doi:10.1007/s00401-012-1057-6

3. Goldfarb LG, Olivé M, Vicart P, Goebel HH. Intermediate Filament Diseases : Desminopathy. 2009:131–164.

4. Piñol-Ripoll G, Shatunov A, Cabello A, et al. Severe infantile-onset cardiomyopathy associated with a homozygous deletion in desmin. Neuromuscul Disord. 2009. doi:10.1016/j.nmd.2009.04.004

5. Palmio J, Penttilä S, Huovinen S, Haapasalo H, Udd B. An unusual phenotype of late-onset desminopathy. Neuromuscul Disord. 2013;23(11):922–923. doi:10.1016/J.NMD.2013.06.374

6. Ky VS, L VH, Jdh J, Walle D. Desmin-related myopathy. 2011:354–366. doi:10.1111/j.1399-0004.2010.01512.x

7. Munoz-Marmol AM, Strasser G, Isamat M, et al. A dysfunctional desmin mutation in a patient with severe generalized myopathy. Proc Natl Acad Sci. 2002. doi:10.1073/pnas.95.19.11312

8. Walter MC, Reilich P, Huebner A, et al. Scapuloperoneal syndrome type Kaeser and a wide phenotypic spectrum of adult-onset, dominant myopathies are associated with the desmin mutation R350P. Brain. 2007. doi:10.1093/brain/awm039

9. Zhang J, Kumar A, Stalker HJ, et al. Clinical and molecular studies of a large family with desmin-associated restrictive cardiomyopathy: Familial restrictive cardiomyopathy. Clin Genet. 2001. doi:10.1034/j.1399-0004.2001.590406.x

10. Clemen CS, Stöckigt F, Strucksberg KH, et al. The toxic effect of R350P mutant desmin in striated muscle of man and mouse. Acta Neuropathol. 2015. doi:10.1007/s00401-014-1363-2

11. Schröder R, Goudeau B, Simon MC, et al. On noxious desmin: Functional effects of a novel heterozygous desmin insertion mutation on the extrasarcomeric desmin cytoskeleton and mitochondria. Hum Mol Genet. 2003. doi:10.1093/hmg/ddg060

12. Herrmann H, Fouquet B, Franke WW. Expression of intermediate filament proteins during development of Xenopus laevis. II. Identification and molecular characterization of desmin. Development. 1989;105(2):299–307.

13. Kaufman SJ, Foster RF. Replicating myoblasts express a muscle-specific phenotype. Proc Natl Acad Sci U S A. 1988;85(24):9606–9610. doi:10.1073/pnas.85.24.9606

14. Schaart G, Viebahn C, Langmann W, Ramaekers F. Desmin and titin expression in early postimplantation mouse embryos. Development. 1989;107(3):585–596.

15. Maddison P, Damian MS, Sewry C, et al. Clinical and myopathological characteristics of desminopathy caused by a mutation in desmin tail domain. Eur Neurol. 2012. doi:10.1159/000341617

16. Su H, Wang X. The ubiquitin-proteasome system in cardiac proteinopathy : a quality control perspective. 2010:253-262. doi:10.1093/cvr/cvp287

17. Cabet E, Batonnet-Pichon S, Delort F, Gausserès B, Vicart P, Lilienbaum A. Antioxidant treatment and induction of autophagy cooperate to reduce desmin aggregation in a cellular model of desminopathy. PLoS One. 2015. doi:10.1371/journal.pone.0137009

18. Goebel HH, Bornemann A. Desmin pathology in neuromuscular diseases. Virchows Arch B. 1993. doi:10.1007/BF02915105

19. Fidziańska A, Ryniewicz B, Barcikowska M, Goebel HH. A new familial congenital myopathy in children with desmin and dystrophin reacting plaques. J Neurol Sci. 1995. doi:10.1016/0022-510X(95)00090-O

20. Goebel HH, Blaschek A. Protein Aggregation in Congenital Myopathies. Semin Pediatr Neurol. 2011. doi:10.1016/j.spen.2011.10.009

21. Bastian A, Goebel HH. Protein aggregation in inclusion body myositis, a sporadic form among protein aggregate myopathies, and in myofibrillar myopathies--a comparative study. Rom J Intern Med. 2010.

22. Goebel HH, Voit T, Warlo I, Jacobs K, Johannsen U, Muller CR. Immunohistologic and electron microscopic abnormalities of desmin and dystrophin in familial cardiomyopathy and myopathy. Rev Neurol. 1994.

23. Vrabie A, Goldfarb LG, Shatunov A, et al. The enlarging spectrum of desminopathies: New morphological findings, eastward geographic spread, novel exon 3 desmin mutation. Acta Neuropathol. 2005. doi:10.1007/s00401-005-0980-1

24. Dagvadorj A, Goudeau B, Hilton-Jones D, et al. Respiratory insufficiency in desminopathy patients caused by introduction of proline residues in desmin C-terminal α-helical segment. Muscle and Nerve. 2003. doi:10.1002/mus.10370

25. Bär H, Fischer D, Goudeau B, et al. Pathogenic effects of a novel heterozygous R350P desmin mutation on the assembly of desmin intermediate filaments in vivo and in vitro. Hum Mol Genet. 2005. doi:10.1093/hmg/ddi136

26. Langer HT, Mossakowski AA, Willis BJ, et al. Generation of desminopathy in rats using CRISPR - Cas 9. 2020;(September):1364–1376. doi:10.1002/jcsm.12619

27. Langer HT, Mossakowski AA, Avey AM, et al. A mutation in desmin makes skeletal muscle less vulnerable to acute muscle damage after eccentric loading in rats | Enhanced Reader. 2021.

28. Holmes WE, Angel TE, Li KW, Hellerstein MK. Dynamic Proteomics: In Vivo Proteome-Wide Measurement of Protein Kinetics Using Metabolic Labeling. In: Methods in Enzymology. Vol 561. Academic Press Inc.; 2015:219–276. doi:10.1016/bs.mie.2015.05.018

29. Armstrong RB, Ogilvie RW, Ogilvie W. Eccentric exercise-induced to rat skeletal muscle. 2021:80–93.

30. Duan C, Delp MD, Hayes DA, et al. Rat skeletal muscle mitochondrial and injury from downhill walking. 2018;(2):1241–1251.

31. Ogilvie RW, Armstrong RB, Baird KE, Bottoms CL. Lesions in the Rat Soleus Muscle Following Eccentrically Biased Exercise. 1988;346.

32. Armstrong RB, Laughlin MH. Metabolism of rats running up and down an incline. 2018:518–521.

33. Armstrong RB, Taylor CR. GLYCOGEN LOSS IN RAT MUSCLES DURING LOCOMOTION ON DIFFERENT INCLINES. 1993;144:135–144.

34. Delp MD, Duan C, Ray CA, et al. Rat hindlimb muscle blood flow during level and downhill locomotion. 2021:564–568.

35. Qin F, Dong Y, Wang S, Xu M, Wang Z, Qu C. Maximum oxygen consumption and quantification of exercise intensity in untrained male Wistar rats. 2020:1–8.

36. Ariza A, Coll J, Fernandez-Figueras MT, et al. Desmin myopathy: a multisystem disorder involving skeletal, cardiac, and smooth muscle. Hum Pathol. 1995;26(9):1032–1037. doi:10.1016/0046-8177(95)90095-0

37. Hughes DC, Marcotte GR, Baehr LM, et al. Alterations in the muscle force transfer apparatus in aged rats during unloading and reloading: impact of microRNA-31. J Physiol. 2018;596(14):2883–2900. doi:10.1113/JP275833

38. Vafiadaki E, Arvanitis DA, Sanoudou D. Muscle LIM Protein: Master regulator of cardiac and skeletal muscle functions. Gene. 2015;566(1):1–7. doi:10.1016/j.gene.2015.04.077

39. Malm C, Yu JG. Exercise-induced muscle damage and inflammation: Re-evaluation by proteomics. Histochem Cell Biol. 2012;138(1):89–99. doi:10.1007/s00418-012-0946-z

40. Roostalu U, Strähle U. In Vivo Imaging of Molecular Interactions at Damaged Sarcolemma. Dev Cell. 2012. doi:10.1016/j.devcel.2011.12.008

41. Bansal D, Miyake K, Vogel SS, et al. Defective membrane repair in dysferlin-deficient muscular dystrophy. Nature. 2003. doi:10.1038/nature01573

42. Glover L, Brown RH. Dysferlin in membrane trafficking and patch repair. Traffic. 2007. doi:10.1111/j.1600-0854.2007.00573.x

43. Bonakdar N, Luczak J, Lautscham L, et al. Biomechanical characterization of a desminopathy in primary human myoblasts. Biochem Biophys Res Commun. 2012;419(4):703–707. doi:10.1016/j.bbrc.2012.02.083

44. Herrmann H, Bär H, Kreplak L, Strelkov S V., Aebi U. Intermediate filaments: From cell architecture to nanomechanics. Nat Rev Mol Cell Biol. 2007;8(7):562–573. doi:10.1038/nrm2197

45. Sam M, Shah S, Fridén J, Milner DJ, Capetanaki Y, Lieber RL. Desmin knockout muscles generate lower stress and are less vulnerable to injury compared with wild-type muscles. Am J Physiol Physiol. 2000;279(4):C1116–C1122. doi:10.1152/ajpcell.2000.279.4.C1116

46. Barash IA, Peters D, Fridén J, Lutz GJ, Lieber RL. Desmin cytoskeletal modifications after a bout of eccentric exercise in the rat. Am J Physiol - Regul Integr Comp Physiol. 2002;283(4 52-4). doi:10.1152/ajpregu.00185.2002

47. Lieber RL, Shah S, Fridén J. Cytoskeletal disruption after eccentric contraction-induced muscle injury. In: Clinical Orthopaedics and Related Research.; 2002. doi:10.1097/00003086-200210001-00011

48. Segard B-D, Delort F, Bailleux V, et al. N-Acetyl-L-Cysteine Prevents Stress-Induced Desmin Aggregation in Cellular Models of Desminopathy. Kampinga HH, ed. PLoS One. 2013;8(10):e76361. doi:10.1371/journal.pone.0076361

49. Totsuka M, Nakaji S, Suzuki K, Sugawara K, Sato K. Break point of serum creatine kinase release after endurance exercise. J Appl Physiol. 2002;93(4):1280–1286. doi:10.1152/japplphysiol.01270.2001

50. Magal M, Dumke CL, Urbiztondo ZG, et al. Relationship between serum creatine kinase activity following exercise-induced muscle damage and muscle fibre composition. J Sports Sci. 2010;28(3):257–266. doi:10.1080/02640410903440892

51. Mohaupt MG, Karas RH, Babiychuk EB, et al. Association between statin-associated myopathy and skeletal muscle damage. CMAJ. 2009;181(1-2):E11. doi:10.1503/cmaj.081785

52. Van Spaendonck-Zwarts KY, Van Hessem L, Jongbloed JDH, et al. Desmin-related myopathy. Clin Genet. 2011. doi:10.1111/j.1399-0004.2010.01512.x

53. Bach M, Larance M, James DE, Ramm G. The serine/threonine kinase ULK1 is a target of multiple phosphorylation events. Biochem J. 2011. doi:10.1042/BJ20101894

54. Kim J, Kundu M, Viollet B, Guan KL. AMPK and mTOR regulate autophagy through direct phosphorylation of Ulk1. Nat Cell Biol. 2011. doi:10.1038/ncb2152

55. Wong PM, Puente C, Ganley IG, Jiang X. The ULK1 complex sensing nutrient signals for autophagy activation. Autophagy. 2013. doi:10.4161/auto.23323

56. Pagano AF, Py G, Bernardi H, Candau RB, Sanchez AMJ. Autophagy and protein turnover signaling in slow-twitch muscle during exercise. Med Sci Sports Exerc. 2014. doi:10.1249/MSS.0000000000000237

57. Møller AB, Vendelbo MH, Christensen B, et al. Physical exercise increases autophagic signaling through ULK1 in human skeletal muscle. J Appl Physiol. 2015. doi:10.1152/japplphysiol.01116.2014

58. Kwon I, Jang Y, Cho JY, Jang YC, Lee Y. Long-term resistance exercise-induced muscular hypertrophy is associated with autophagy modulation in rats. J Physiol Sci. 2018. doi:10.1007/s12576-017-0531-2

59. Ogura Y, Iemitsu M, Naito H, et al. Single bout of running exercise changes LC3-II expression in rat cardiac muscle. Biochem Biophys Res Commun. 2011. doi:10.1016/j.bbrc.2011.09.152

60. Phillips BE, Williams JP, Gustafsson T, et al. Molecular Networks of Human Muscle Adaptation to Exercise and Age. PLoS Genet. 2013. doi:10.1371/journal.pgen.1003389

61. Bajor M, Zych AO, Graczyk-Jarzynka A, et al. Targeting peroxiredoxin 1 impairs growth of breast cancer cells and potently sensitises these cells to prooxidant agents. Br J Cancer. 2018. doi:10.1038/s41416-018-0263-y

62. Kim Y, Jang HH. The Role of Peroxiredoxin Family in Cancer Signaling. J Cancer Prev. 2019. doi:10.15430/jcp.2019.24.2.65

63. Langer HT, Afzal S, Kempa S, Spuler S. Nerve damage induced skeletal muscle atrophy is associated with increased accumulation of intramuscular glucose and polyol pathway intermediates. Sci Rep. 2020;10(1). doi:10.1038/s41598-020-58213-1

64. Bostock EL, Edwards BT, Jacques MF, et al. Impaired glucose tolerance in adults with duchenne and becker muscular dystrophy. Nutrients. 2018. doi:10.3390/nu10121947

65. Schneider SM, Sridhar V, Bettis AK, et al. Glucose Metabolism as a Pre-clinical Biomarker for the Golden Retriever Model of Duchenne Muscular Dystrophy. Mol Imaging Biol. 2018. doi:10.1007/s11307-018-1174-2

66. Xiao Y, Zhu H, Li L, et al. Global analysis of protein expression in muscle tissues of dermatomyositis/polymyosisits patients demonstrated an association between dysferlin and human leucocyte antigen A. Rheumatol (United Kingdom). 2019. doi:10.1093/rheumatology/kez085

67. Mille-Hamard L, Billat VL, Henry E, et al. Skeletal muscle alterations and exercise performance decrease in erythropoietin-deficient mice: A comparative study. BMC Med Genomics. 2012;5(March 2014). doi:10.1186/1755-8794-5-29

68. Gasier HG, Riechman SE, Wiggs MP, Previs SF, Fluckey JD. A comparison of 2H2O and phenylalanine flooding dose to investigate muscle protein synthesis with acute exercise in rats. Am J Physiol - Endocrinol Metab. 2009;297(1). doi:10.1152/ajpendo.90872.2008

69. Busch R, Kim YK, Neese RA, et al. Measurement of protein turnover rates by heavy water labeling of nonessential amino acids. Biochim Biophys Acta - Gen Subj. 2006;1760(5):730–744. doi:10.1016/j.bbagen.2005.12.023

70. Woessner JF. The determination of hydroxyproline in tissue and protein samples containing small proportions of this imino acid. Arch Biochem Biophys. 1961;93(2):440–447. doi:10.1016/0003-9861(61)90291-0

71. Creemers LB, Jansen DC, Van Veen-Reurings A, Van Den Bos T, Everts V. Microassay for the assessment of low levels of hydroxyproline. Biotechniques. 1997;22(4):656–658. doi:10.2144/97224bm19

72. Yang D, Diraison F, Beylot M, et al. Assay of low deuterium enrichment of water by isotopic exchange with [U-13C3]acetone and gas chromatography-mass spectrometry. Anal Biochem. 1998;258(2):315–321. doi:10.1006/abio.1998.2632

73. Huang DW, Sherman BT, Lempicki RA. Systematic and integrative analysis of large gene lists using DAVID bioinformatics resources. Nat Protoc. 2009;4(1):44–57. doi:10.1038/nprot.2008.211

74. Hedges L V. Distribution Theory for Glass’s Estimator of Effect Size and Related Estimators. J Educ Stat. 1981;6(2):107–128. doi:10.2307/1164588

